# Studies of human twins reveal genetic variation that affects dietary fat perception

**DOI:** 10.1101/2020.01.18.910448

**Authors:** Cailu Lin, Lauren Colquitt, Paul Wise, Paul A. S. Breslin, Nancy E. Rawson, Federica Genovese, Ivy Maina, Paule Joseph, Lydia Fomuso, Louise Slade, Dennis Brooks, Aurélie Miclo, John E. Hayes, Antonio Sullo, Danielle R. Reed

## Abstract

To learn more about the mechanisms of human dietary fat perception, 398 human twins rated fattiness and liking for six types of potato chips that differed in triglyceride content (2.5, 5, 10, and 15% corn oil); reliability estimates were obtained from a subset (*n* = 50) who did the task twice. Some chips also had a saturated long-chain fatty acid (hexadecanoic acid, 16:0) added (0.2%) to evaluate its effect on fattiness and liking. We computed the heritability of these measures and conducted a genome-wide association study (GWAS) to identify regions of the genome that co-segregate with fattiness and liking. Perceived fattiness and liking for the potato chips were reliable (*r* = 0.31-0.62, *p* < 0.05) and heritable (up to *h*^2^ = 0.29, *p* < 0.001, for liking). Adding hexadecanoic acid to the potato chips significantly increased ratings of fattiness but decreased liking. Twins with the G allele of *rs263429* near *GATA3-AS1* or the G allele of *rs8103990* within *ZNF729* reported more liking for potato chips than did twins with the other allele (multivariate GWAS, *p* < 1×10^-5^), with results reaching genome-wide suggestive but not significance criteria. Person-to-person variation in the perception and liking of dietary fat was (a) negatively affected by the addition of a saturated fatty acid and (b) related to inborn genetic variants. These data suggest liking for dietary fat is not due solely to fatty acid content and highlight new candidate genes and proteins within this sensory pathway.

## Introduction

Sensory nutrition is a research area that investigates how the taste, smell, and flavor of food and drink affect food intake and diet quality, and how food choice in turn affects human health and disease (Forde 2018; Hayes 2015). While food is essential to our survival, and eating may be pleasant, it can also be dangerous, especially for those who “dig their grave with a spoon” (Card 2013) and die from heart disease or diabetes, health conditions that arise in whole or in part from dietary choices (Reed and Knaapila 2010). Some of the pleasure of food arises from its dietary fat and sugar content. The sweetness of sugar is well understood from a sensory perspective (Nelson *et al.* 2001), with direct links between taste cells and brain areas of reward (e.g., (Veldhuizen *et al.* 2017)). In contrast, the initial sensory steps responsible for the perception of dietary fat are less well understood, and what is known is contentious: whether there is a distinct taste quality for fat or fatty acids, and which of the chemical and texture components of fat are responsible for the sensations it evokes (Reed and Xia 2015; Running *et al.* 2015; Running and Mattes 2016).

One unresolved conundrum is mounting evidence that, while triglycerides and fatty acids both impart fatty sensations in foods, triglycerides tend to have a positive hedonic valance, e.g.,(Bakke *et al.* 2016) whereas fatty acids typically have a negative hedonic valence, e.g., scratchy (Voigt *et al.* 2014) or otherwise “bad” (Running and Mattes 2016). These data suggest multiple sensory pathways are involved in the perception of fats in foods (Drewnowski 1992). One method to learn more about these multiple pathways is to evaluate origins of person-to-person or animal-to-animal differences—this type of genetics-driven approach helped identify the bitter and sweet receptors (Reed and Knaapila 2010; Reed *et al.* 2006). Here, we reasoned that people differ in their response to fat in food, that these differences are heritable, and that genome-wide methods are likely useful to identify the relevant genes.

To establish heritability, we selected a classic twin design, comparing monozygotic (MZ) and dizygotic (DZ) twins for their response to fat in foods. We also had to choose appropriate test stimuli that would generalize to real foods (vs. model systems) and appropriate behavioral methods. No one standard method has been adopted, with investigators in this area using many different stimuli to measure fat perception, including oil-and-water mixtures (Heinze *et al.* 2017); oil in salad dressing (Keller *et al.* 2012); fat in puddings (Mennella *et al.* 2012), in scrambled eggs or mashed potatoes (Mela and Sacchetti 1991), or in ice cream (Rolon *et al.* 2017) or added fatty acids in chocolate (Running *et al.* 2017). Here we used potato chips that varied in amounts of corn oil and an added fatty acid, capitalizing on our technical expertise in their production and practical constraints of our testing environment (an annual convention of twins; see below). We also tested the twins’ ability to discriminate high- and low-fat milk samples.

## Materials and Methods

### Participants

We tested adult MZ and DZ twins who attended an annual convention of twins, the Twin Days Festival in Twinsburg, OH. This event is held each August, and all data reported here were collected during the 2018 convention. The exclusion criteria for participation were age less than 18 years, pregnancy, or an allergy or sensitivity to milk. All data were collected under protocols approved by the University of Pennsylvania Institutional Review Board (#701426).

### Stimuli

Three types of stimuli were used: potato chips that differed in triglyceride and fatty acid content, multiple prototypical tastants, and milk that was either high (18.00%) or low (2.35%) in fat. Six types of potato chips were prepared, following standard methods at Pepsico research laboratories: chips that contained 2.5, 5.0, 10, or 15% corn oil and chips with 2.5% or 5.0% corn oil with added 0.2% (w/w) hexadecanoic acid, a saturated long-chain fatty acid (16:0). Time constraints prevented us from testing all combinations of triglycerides and fatty acids. Ascending amounts of corn oil were chosen to minimize carryover effects across samples; the fatty acid was added to gauge its impact on ratings of fattiness and liking.

The second type of stimuli comprised standard solutions (5 mL) used in taste psychophysics: plain deionized water, sucrose (12% w/v, 350 mM), sodium chloride (1.5% w/v, 256 mM), and the bitter compound phenylthiocarbamide (PTC; 1.8 ×10^-4^ M), all purchased from Sigma (St. Louis, MO). (We also tested menthol [1 mM] and capsaicin [3 μM] for an unrelated project; those results are not reported here.) The third stimuli comprised milk with 18.00% or 2.35% fat mixed at the Monell Chemical Senses Center using Shop Rite brand instant nonfat dry milk (SKU/UPC 041190010189) purchased at a local grocery store and anhydrous dairy fat (**Table 1**). All ingredients were combined in a homogenizer (GEA, Düsseldorf, Germany) and processed with five passes at 250 bars of pressure; resulting particle sizes were within the expected range.

**Table 1.** High- and low-fat milk ingredients

| Milk Type | Fat Content<br>(%) | Water<br>(mL) | Dry Milk<br>(g) | Dairy Fat<br>(g) | Casein<br>(g) |
| --- | --- | --- | --- | --- | --- |
| Low fat | 2.35 | 890 | 90.7 | 23.7 | 10.09 |
| High fat | 18.00 | 890 | 90.7 | 216.8 | 12.04 |

### Sample presentation

Single potato chips of roughly equivalent size and weight were placed in clear 3-5 oz plastic souffle cups with plastic lids (Universal Product Code [UPC] #742010492467). Participants were given potato chips in a predetermined order and asked to rate the potato chips for “fattiness” and “liking” on visual analog scales presented on an Apple iPad Air (9.7-inch display; Apple Inc., Cupertino, CA). Liking scales were anchored with “do not like at all” on the left and “like extremely” on the right. Similarly, the fattiness scale was anchored on the left with “not fatty at all” and on the right with “extremely fatty.” We also asked about “crispiness” and “saltiness,” to prevent a halo-dumping effect, a bias in sensory ratings which can occur when subjects are provided too few salient rating options (Clark and Lawless 1994). Participants were instructed to rinse their mouth with water (Nestle Pure Life, UPC 068274934711) after each sample. For logistical reasons and enhanced ecological validity, participants did not wear nose clips and chewed and swallowed all potato chip samples.

For the taste solutions, participants rated each for the qualities of “liking,” “saltiness,” “sweetness,” “sourness,” “bitterness,” and “burn” on visual analog scales, with the left side anchored with “no [quality] at all” and the right side anchored with “extreme [quality],” as previously described (Knaapila *et al.* 2012). To focus on taste and reduce odor cues, participants wore nose clips (GENEXA LLC, UPC 708981350007). Participants were asked to hold each solution in their mouth for 5 s, rate it on the scale provided, spit out the solution, and rinse their mouth with water afterward.

For the milk fat discrimination test, a two-alternative forced choice task was used. Before testing began, each participant was given two references as warm-up samples; these were verbally identified to participants as “low-fat” and “high-fat” samples, respectively. Participants were then given 10 pairs of opaque bottles (EP-34434, Berry Global Group, Inc.). Each pair contained one low-fat and one high-fat sample (each 5 mL) presented in a fixed order. Participants wore nose clips; they were instructed to hold each sample in their mouth for 5 s, spit out the sample, and rinse their mouth with water afterward. For each pair, participants were asked, “Which solution tastes fattier?” If they were unsure, they were instructed to guess. Discrimination ability was defined as the number correct across all 10 trials (i.e., perfect discrimination would be 10 out of 10 trials correct).

### Saliva collection and DNA extraction

We obtained saliva samples from all participants by asking them to expectorate into collection tubes; DNA was extracted from the saliva using procedures recommended by Oragene (DNA Genotek, Kanata, Canada). We measured and recorded DNA concentration and quality scores using a Nanodrop 1000 Spectrophotometer (Thermo Fisher Scientific, Waltham, MA).

### Genotyping

We conducted both single-marker and high-throughput based genotyping. Using the single-marker method, we typed three variant sites in the *TAS2R38* gene in all twins, as a quality-control step (a) to ensure that the DNA extracted from saliva could be genotyped, (b) to confirm that the genotype matched the psychophysical ratings of PTC bitterness, and (c) to get preliminary confirmation of twin zygosity (each pair of MZ twins is expected to have the same genotype). For these assays, DNA samples were diluted to a concentration of 10 ng/μL and used as templates in Taqman assays (*rs713598*, C__8876467_10; *rs1726866*, C__9506827_10; and *rs10246939*, C__9506826_10; Applied Biosystems, Foster City, CA) using previously established methods.

For the DNA high-throughput genotyping, we sent the DNA samples to the Center for Inherited Disease Research (CIDR; Baltimore, MD), which typed them for the Illumina OmniExpress panel (Infinium OmniExpressExome-8, v1.6; Illumina, San Diego, CA) following the manufacturer’s procedures and the CIDR’s standard quality-control methods. For 176 MZ twin pairs, we used high-throughput genotyping for only one twin of each pair and imputed the genotype of the other member of the pair because of their presumed identical genomes.

### Twin zygosity

Twin zygosity was measured in three ways. Twins self-reported their zygosity status as (a) monozygotic (MZ; identical), (b) dizygotic (DZ; fraternal), or (c) uncertain; photographs were taken of each twin and rated for physical similarity by a research assistant blind to self-reported zygosity, and all twins were genotyped for the three markers described above. In rare cases were zygosity status was still uncertain, both members of the pair were genotyped using the high-throughput-based genotyping method (see above).

### Data analysis

We conducted four types of statistical analysis: (a) descriptive statistics of the psychophysical data, (b) calculation of heritability, (c) tests of genome-wide association between genetic variants and the measures of fat perception, and (d) gene expression (RNASeq) and bioinformatics (enrichment) analyses. All descriptive statistics, such as means, standard deviations (SDs), and correlations among variables, were computed using R (v. 3.53) and R-Studio (v. 1.1.456).

#### Sensory analyses

For descriptive analyses, we plotted the probability density of the data (smoothed by a kernel density estimator) by a violin plot, calculated mean and SD, and checked for sex, race, and age effects on the sensory measures in a general linear model (GLM) using race and sex as fixed effects and age as a covariate. For all GLM analyses, individual group means were evaluated for difference using Tukey post hoc tests (honestly significant difference [HSD]). If race and sex had a significant effect in the GLM analysis, to better understand their effects on psychophysical outcomes we grouped participants by these factors and compared the mean ratings. For age and its relationship to the psychophysical measures, we computed Pearson correlations.

To evaluate whether there were consistent person-to-person differences in the rating of the potato chips overall, Pearson correlations of intensity and liking measures among the six types of potato chip were calculated. In addition, we calculated Cronbach’s alpha for psychophysical measures across all six types of potato chips. To understand the reliability of the measures, we assessed test-retest correlations among the same measures taken twice in a subset of participants (n = 50).

To gauge the effect of corn oil concentrations and hexadecanoic acid on the sensory measures, we reconducted a linear mixed-model analysis with corn oil concentration (2.5% and 5.0%) and hexadecanoic acid (added or not) as two separate factors and treated the psychophysical data as repeated measurements, with race and age as covariates in the model. (We did not include sex in this model because results indicated that male and females were similar in their ratings.) In a complementary analysis, we reconducted the analysis using potato chip type as a single factor (with six levels, one for each type of potato chip). These complementary analyses were included because of the unbalanced design: not all concentrations of corn oil were presented with and without the added 0.2% hexadecanoic acid.

#### Heritability

For the heritability analysis, the Cholesky model was used to evaluate the magnitude of genetic and environmental influences on the traits, and the phenotypic variance was decomposed into additive genetic component (a^2^), shared environmental factors (c^2^), and nonshared environmental or individual-specific factors (e^2^), as described previously (Wise *et al.* 2007). Variance accounted for by each of these components was calculated by comparing MZ twin correlations to DZ twin correlations. The computation of the heritability was conducted using R package OpenMx (v. 2.13) (Boker *et al.* 2011).

#### Genome-wide association studies

For GWAS we expanded variants from ∼720,000 to 11,315,231 by imputation using the Michigan Imputation Server (Das *et al.* 2016) with the reference genome HRCr1.1 (McCarthy *et al.* 2016). We filtered out markers with a low minor allele frequency (<5%) and removed markers that had *p*-values associated with Hardy-Weinberg disequilibrium < 1e-6, genotype call rate < 0.9, and imputation score < 0.3. The remaining 4,234,798 variants on the 22 autosomes were used for GWAS for each trait (univariate GWAS [uvGWAS]), with genetic relatedness matrix (20 eigenvalues) calculated by principal components analysis, and sex and age used as covariates (Hwang *et al.* 2019; Liu *et al.* 2018; Wu *et al.* 2018). The genome-wide significance threshold was *p* = 5.0e-8, and for suggestive associations it was *p* = 1e-5 (International HapMap 2005; Pe’er *et al.* 2008).

We reasoned that there would be more statistical power to detect associations if we considered the liking and fattiness ratings from all potato chips simultaneously, especially because, as the results indicated, these measures were correlated (e.g., people with high liking ratings for the 5% corn oil chip also liked the 10% chip more). Thus, we conducted multivariate GWAS (mvGWAS) using the correlated ratings for all the potato chips. The covariates are the same as uvGWAS procedure; the computation was done using GEMMA (Zhou and Stephens 2012), and regional associational plots were created using LocusZoom (Pruim *et al.* 2010). For the mvGWAS, GEMMA adjusted for testing multiple phenotypes and applied a correction for multiple phenotypes (Fatumo *et al.* 2019). For the milk discrimination task, the trait was not heritable (see Results), so we did not conduct GWAS.

#### Candidate gene analyses

We extracted variants from the candidate genes that were previously implicated in the sensory signaling of fat taste from either animal models (mouse and rat) or human studies: *CD36* (Abumrad 2005; Gaillard *et al.* 2007; Keller *et al.* 2012; Laugerette *et al.* 2005; Pepino *et al.* 2012; Sclafani *et al.* 2007a), *GNAT3* (Sclafani *et al.* 2007b), *GPR120* (Cartoni *et al.* 2010; Matsumura *et al.* 2007; Tsuzuki 2007), *GPR40* (Cartoni *et al.* 2007; Cartoni *et al.* 2010; Matsumura *et al.* 2007), *TRPM5* (Liu *et al.* 2011; Sclafani *et al.* 2007b), *GPR41* and *GPR43* (Brown *et al.* 2003), *GPR84* (Wang *et al.* 2006), and *KCNA2* (Gilbertson *et al.* 1998; Liu *et al.* 2005). In addition, we looked at genes for salivary enzymes (lipase, lysozyme, and amylase) and protein (lipocalin, mucin, and protein rich in proline) because these proteins change in response to dietary fat consumption (Feron G 2013; Mounayar *et al.* 2014).

To extract the results of genotype-phenotype association for these candidate genes, we conducted analyses using two methods. In method 1 we identified the most significant variant within each candidate genes for each trait and extracted the relevant *p*-value and other test statistics. In method 2 we chose the most significant variant for traits of the potato chip with 5% corn oil (with no added fatty acid) and examined all the sensory measures for the same variant; that is, we chose the 5% corn oil chip as the baseline from which to compare the other associations. These methods are complementary because method 1 detects associations that are specific to a particular concentration of triglyceride and fatty acid combination, while method 2 detects common variants affecting the intensity and liking measures across the potato chip types. We also examined the effect of the variant *rs1761667* within *CD36* because it was previously associated with fat sensory perception in humans (Keller *et al.* 2012; Mrizak *et al.* 2015; Pepino *et al.* 2012; Sayed *et al.* 2015).

#### Gene expression in human taste tissue using the RNASeq method

To understand whether the genes identified by GWAS might be acting at the level of the receptors in taste tissue (as opposed to in the brain or in other tongue tissue, e.g., the filiform papillae), we compared the mRNA expression of these genes to those previously implicated in the peripheral aspects of fat taste perception (e.g., the candidate gene *CD36*) in human taste tissue. To do so, we collected fungiform papillae from subjects recruited for our previous study (Douglas *et al.* 2019) using published procedures (Spielman *et al.* 2010) and isolated the RNA following the manufacturer’s directions, processing the taste tissue with Quick-RNA MiniPrep R1054 (Zymo Research, Irvine, CA). We evaluated RNA quality expressed as an RNA integrity number (RIN) using the Agilent 2200 TapeStation system (Agilent Technologies, Santa Clara, CA). The six samples with sufficient RNA quality as determined by the Next-Generation Sequencing Core of the University of Pennsylvania (RIN > 7; 5 males and 1 female) were used to perform library preparation and sequencing (100 bp single-end) on the HiSeq 4000 sequencer (Illumina, San Diego, CA) following the manufacturer’s sequencing protocols. We mapped reads to the reference genome (GRCh38.p10) after the raw sequence data in fastq format passed standard quality filters equipped in Trimmomatic (Bolger *et al.* 2014), and then normalized the counts using the R package Ballgown (Frazee *et al.* 2014). The expression level in RPKM (reads per kilobase per million mapped reads) of each gene for each sample was used to compare their expression level.

#### Pathway and gene set enrichment analysis

We reasoned that genes identified through GWAS may be partners with other genes that code for proteins in related sensory pathways. Thus, we conducted pathway analyses of the genes identified by uvGWAS and mvGWAS. Using the background of the genes from the database of Gene Ontology annotations (Thomas *et al.* 2003) and Reactome annotations (Fabregat *et al.* 2018; Fabregat *et al.* 2017), we used Fisher’s exact test to examine whether there was enrichment of these pathways versus all annotated human genes using GENEVESTIGATOR (Hruz *et al.* 2008).

## Results

### Participant characteristics

The twins (*N* = 398) were predominantly female (72%, n =285; and 28% male, n = 113), middle-aged (38.6 ± 16.7, mean ± SD), and members of MZ twin pairs (*n* = 360 twins, 90.4%). Most were of European descent (*n* = 331, 83.2%), but some participants were of African descent (*n* = 50, 12.6%). The remaining racial groups (e.g., Asian) were grouped into an “other” category for the analyses described below (*n* = 19, 4.8%). A total of 213 individual subjects were genotyped using the chip-based platform (MZ, *n* = 184; DZ, *n* = 29), and 176 MZ twins had their genotypes imputed.

### Liking and intensity measures

#### Liking and fattiness ratings differed across potato chips with variable fat content

Overall, participants liked the potato chips and were able to accurately rate them for fattiness. Adding 0.2% hexadecanoic acid to the potato chips increased fattiness at both corn oil concentrations tested (**Figure 1A**). The effect of added hexadecanoic acid on liking was less straightforward: for the 5% corn oil chips, adding a 16:0 fatty acid did not alter liking, while for the 2.5% corn oil chips, adding the fatty acid decreased liking (**Figure 1A**). For chips with no added fatty acid, there was a mostly linear increase in ratings of fattiness as corn oil concentration increased, although a plateau was reached above 10% oil (**Figure 1B**). For liking, there was a J-shaped curve: participants liked the 2.5% and 15% corn oil potato chips best (**Figure 1B**). See **Supplemental Figures 1 and 2** and **Supplemental Table 1** for additional details.

**Figure 1.**
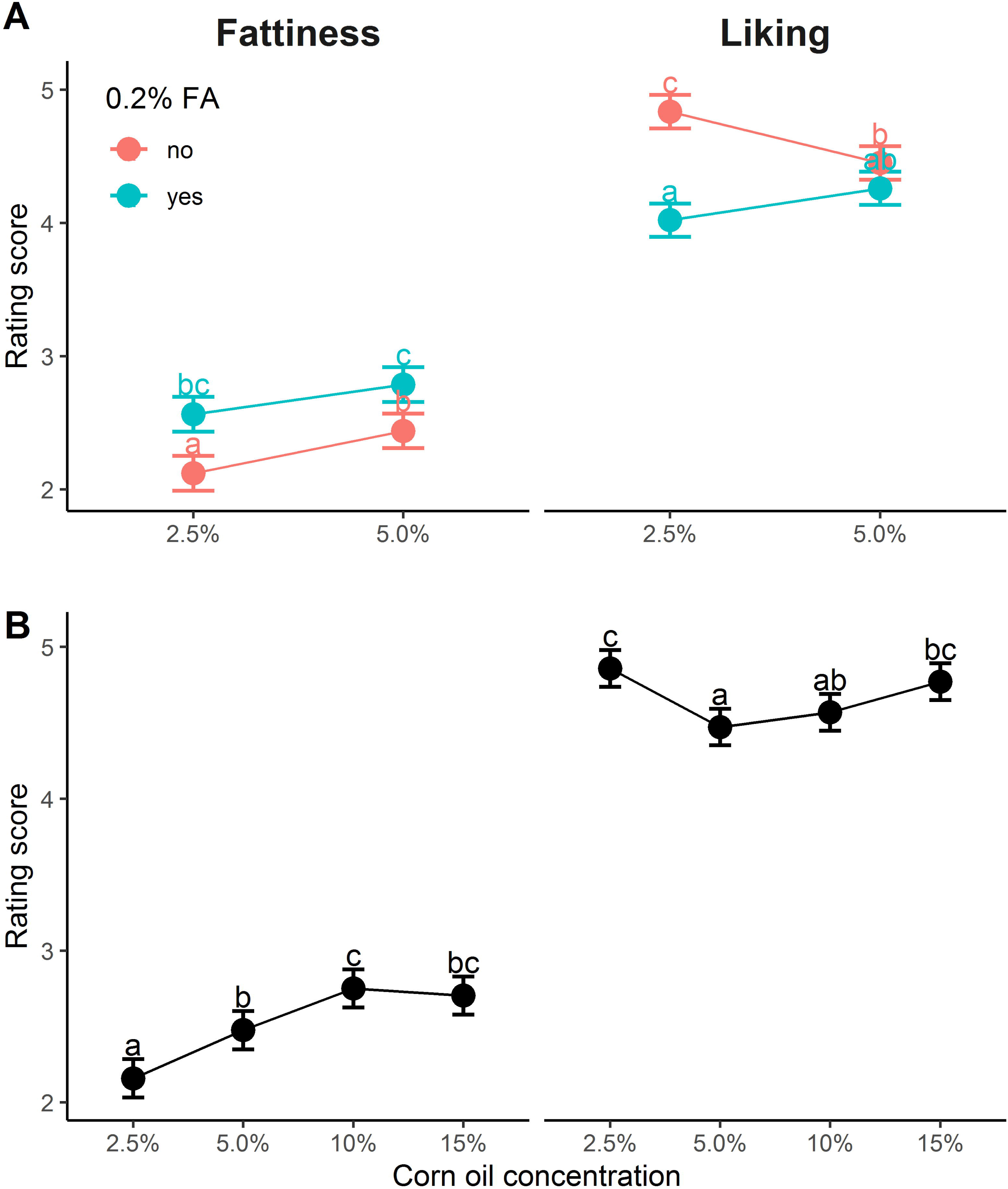
Corn oil and corn oil spiked with 0.2% hexadecanoic acid (FA) modify ratings of fattiness and liking of potato chips. (**A**) Potato chips with more corn oil plus added FA increased fattiness and decreased liking. (**B**) As corn oil concentration (2.5%, 5.0%, 10%, and 15%, without added FA) increased, fattiness ratings increased linearly but liking changed in a J-curve: participants liked potato chips more with corn oil at the lowest and highest concentrations (2.5% and 15%). The points and bars show least square mean (LSM) and standard error of rating scores, and different letters (a, b, and c) indicate a significant LSM difference between groups.

#### Relationship between liking and fattiness relative to benchmarks

Within each type of potato chip, the ratings of liking and fattiness were only slightly or not at all related (**Figure 2A**). This relationship between liking and sensory quality differed from those for the benchmark taste solutions; for example, participants liked sucrose better if they rated it as sweeter (**Figure 2B**).

**Figure 2.**
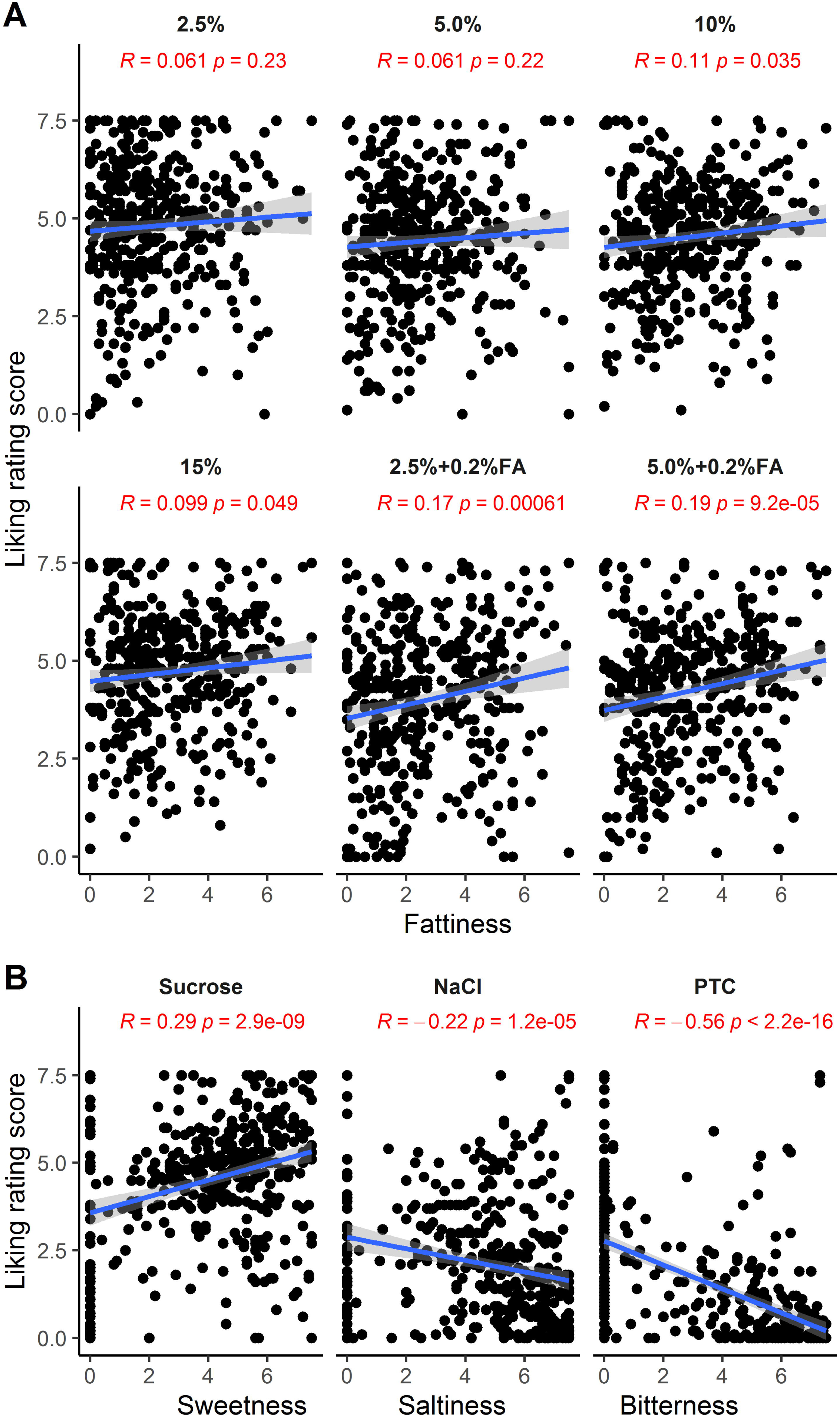
Pearson correlations between sensory measures indicate multiple mechanisms underlying dietary fat perception (*N* = 398). (**A**) No or weak correlations between ratings of liking and fattiness depending on the type of potato chip. FA=fatty acid (hexadecanoic acid). (**B**) Strong correlations between liking and other taste ratings (sweetness, saltiness, and bitterness) for the standard taste solutions sucrose, NaCl, and phenylthiocarbamide (PTC).

#### Reliability of liking and fattiness relative to benchmarks

The ratings of both fattiness and liking for the potato chips were reliable (*r* = 0.31-0.62, *p* < 0.05; **Supplemental Figure 3**), slightly lower than (but mostly similar to) those for the benchmark taste solutions (sucrose, NaCl, and PTC; *r* = 0.54-0.74, *p* < 0.0001, except for NaCl saltiness; **Supplemental Figure 3**).

#### Age, race, and sex effects on fattiness and liking

Men and women were similar in their ratings of all sensory stimuli (**Supplemental Table 2**). Race and age had significant effects on some sensory ratings (*p* < 0.01; **Supplemental Table 2**). Younger participants liked some of the potato chip types more than older participants (*r* = −0.17 to −0.14, *p* < 0.001; **Supplemental Figure 4**). People of European ancestry rated some potato chips as less fatty than did people of African ancestry (5.0% corn oil without added fatty acid; *p* < 0.05, GLM analysis followed by post hoc Tukey HSD tests; **Supplemental Figure 5**). There were also race effects for the other sensory stimuli, for example, for the liking of sucrose and PTC. **Supplemental Figure 5** summarizes all sensory results that differed by race.

#### Relationships of ratings across potato chip type

Each participant tasted and rated six potato chips, and there were correlations among each participant’s ratings of fattiness (Cronbach’s alpha = 0.75, 95% confidence boundaries = 0.72-0.79) and liking (Cronbach’s alpha = 0.77, 95% confidence boundaries = 0.74-0.81). Fattiness correlations tended to be higher among the chips without added FA than with the chips with added FA. A scatter matrix of pairwise correlations between potato chips types is shown in **Figure 3**.

**Figure 3.**
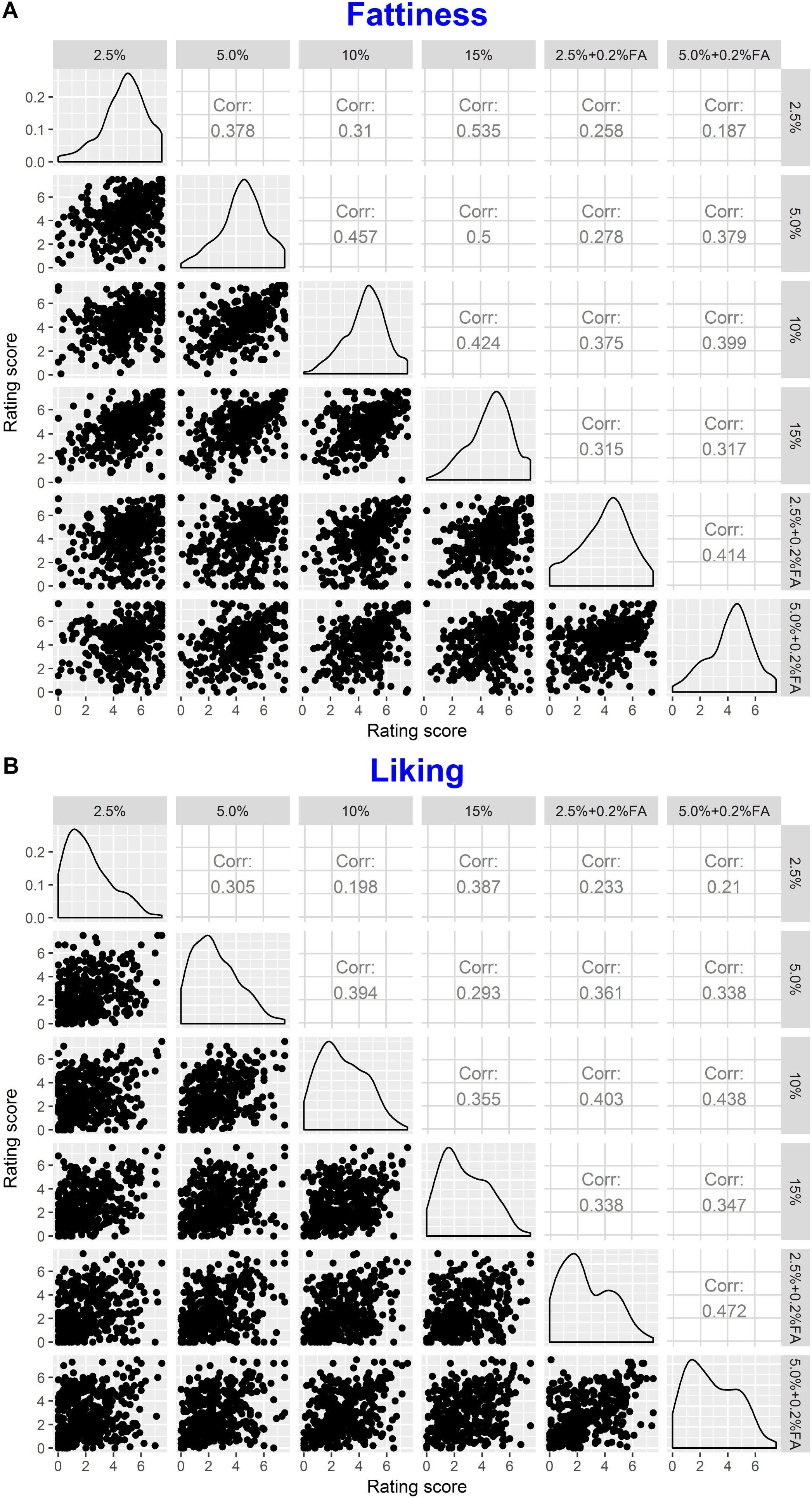
Strong and positive interrelated correlations of ratings of fattiness (A) and liking (B) across the six types of potato chips: scatter plots (lower left), density distributions (diagonal line), and correlations (upper right). FA=fatty acid (hexadecanoic acid).

#### Discrimination of milk fattiness

On average, participants could discriminate the high- and low-fat milk samples (exact binomial test, one-tailed, *p* < 0.0001), but only slightly above chance (probability of success = 0.53; **Supplemental Figure 6A**). This ability to discriminate was only somewhat reliable when testing the same participant twice (retest correlation, *r* = 0.36; *p* > 0.05; **Supplemental Figure 3**). We had expected based on our pilot data collected in our sensory laboratory that about 30% of participants would perform this discrimination perfectly every time, with 10 out of 10 samples correctly identified, but our results showed that only 3% of subjects could do so.

### Heritability

Between about 10% and 30% of the variation in potato chip liking arose from genetics (*h*^2^), but only about 5-15% for ratings of fattiness (**Table 2**). For comparison, for the bitter compound PTC, the most heritable taste trait currently known, liking heritability was 53%, and for sucrose, which has a midrange heritability, it was 46%. The pattern of heritability for NaCl was similar to that for potato chips, as rating of NaCl liking has more genetic variation than does rating of NaCl saltiness. We did not calculate heritability for the milk fat discrimination because there was no similarity in milk discrimination scores between the twins (**Supplemental Figure 6B**).

**Table 2.** Heritability (*h*^2^) of fat sensory traits, with NaCl, sucrose, and PTC as a benchmarks (*n* = 199 twin pairs)

| Stimulus | $h^2$ | CI |
| --- | --- | --- |
| Liking (chips) |  |  |
| 2.5% corn oil | 0.21* | 0.07 – 0.34 |
| 5.0% corn oil | 0.10 | 0.00 – 0.24 |
| 10% corn oil | 0.10 | 0.00 – 0.24 |
| 15% corn oil | 0.29* | 0.15 – 0.41 |
| 2.5% corn oil with 0.2% hexadecanoic acid | 0.10 | 0.00 – 0.24 |
| 5.0% corn oil with 0.2% hexadecanoic acid | 0.10 | 0.00 – 0.24 |
| Fattiness |  |  |
| 2.5% corn oil | 0.05 | 0.00 – 0.20 |
| 5.0% corn oil | 0.11 | 0.00 – 0.25 |
| 10% corn oil | 0.12 | 0.00 – 0.27 |
| 15% corn oil | 0.07 | 0.00 – 0.22 |
| 2.5% corn oil with 0.2% hexadecanoic acid | 0.03 | 0.00 – 0.17 |
| 5.0% corn oil with 0.2% hexadecanoic acid | 0.15 | 0.00 – 0.29 |
| Other solutions |  |  |
| Sucrose sweetness | 0.11 | 0.00 – 0.25 |
| Sucrose liking | 0.46* | 0.33 – 0.56 |
| NaCl saltiness | 0.19* | 0.05 – 0.32 |
| NaCl liking | 0.38* | 0.25 – 0.49 |
| PTC bitterness | 0.49* | 0.38 – 0.59 |
| PTC liking | 0.53* | 0.42 – 0.62 |

CI=confidence interval.
\*Different from zero.

### Genome-wide association

No associations met the commonly accepted genome-wide significance threshold, but we did identify suggestive variants using the univariate and multivariate methods. uvGWAS identified nine associations for fattiness and eight for liking (**Table 3**). All these associations were specific for potato chip type. The mvGWAS detected two variants for chip fattiness and five variants for chip liking (**Table 4**). We reasoned that associations detected with both uvGWAS and mvGWAS would be most valid. Of the seven genotype associations detected by mvGWAS, two (*GATA3-AS1* and *ZNF729)* were also detected by uvGWAS (**Figure 4**): twins with the G allele of *rs263429* (*10:8085050*, near *GATA3-AS1*) reported more liking for the potato chips than did twins with the other allele and the same was true for the G allele of *rs8103990 (19:22476027*, within *ZNF729)* (mvGWAS, *p* < 1×10^-5^; **Table 4**). We show the allelic effects for these two variants in **Figure 5**. The effects of the novel variants were larger than those for *CD36*, the candidate gene previously associated with fat perception (**Supplemental Figure 7**).

**Figure 4.**
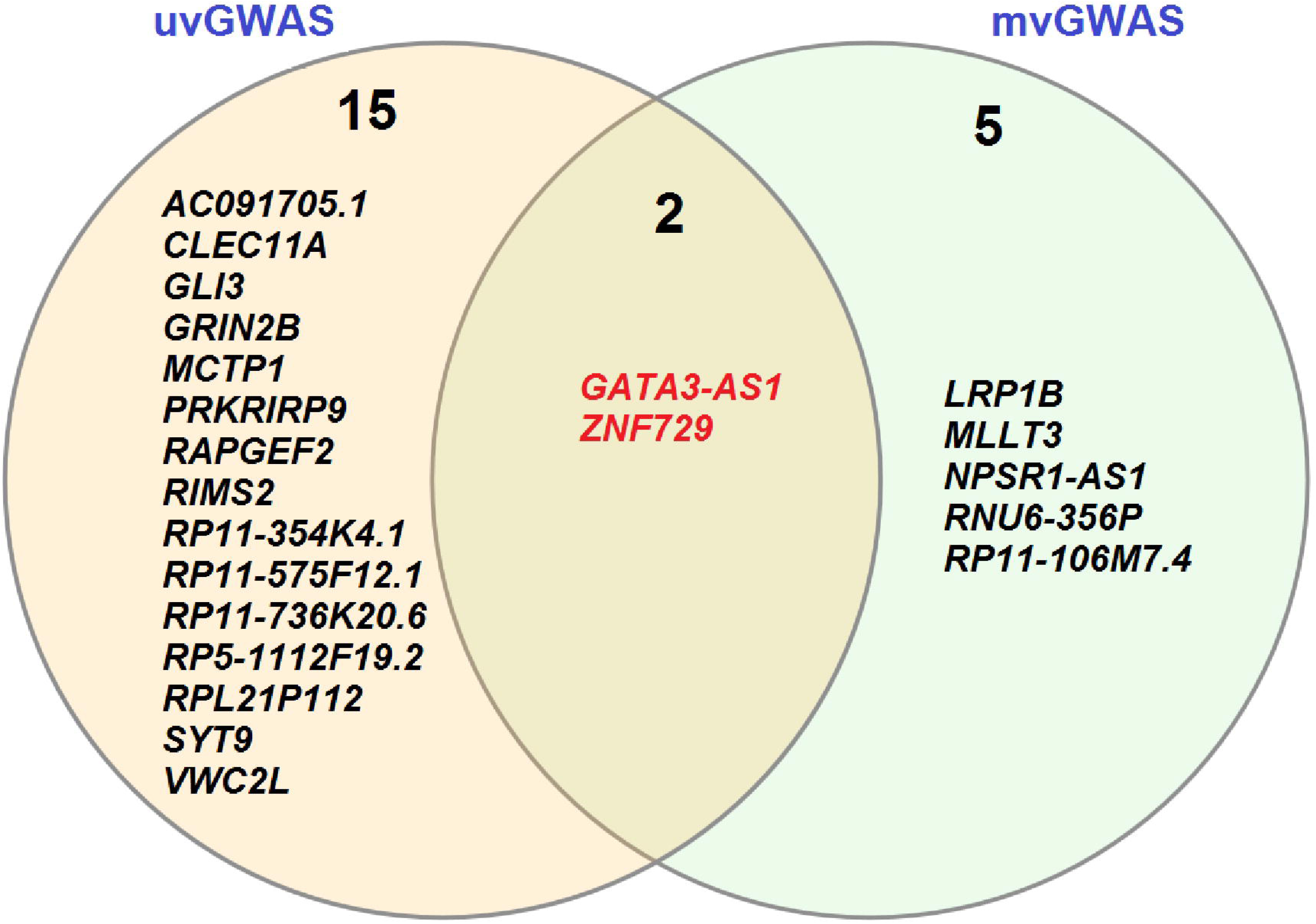
Venn diagram comparing loci identified by uvGWAS and mvGWAS (see Methods for details). Two variants were detected by both methods: *10:8085050* near the gene *GATA3-AS1* and *19:22476027* within *ZNF729*.

**Figure 5.**
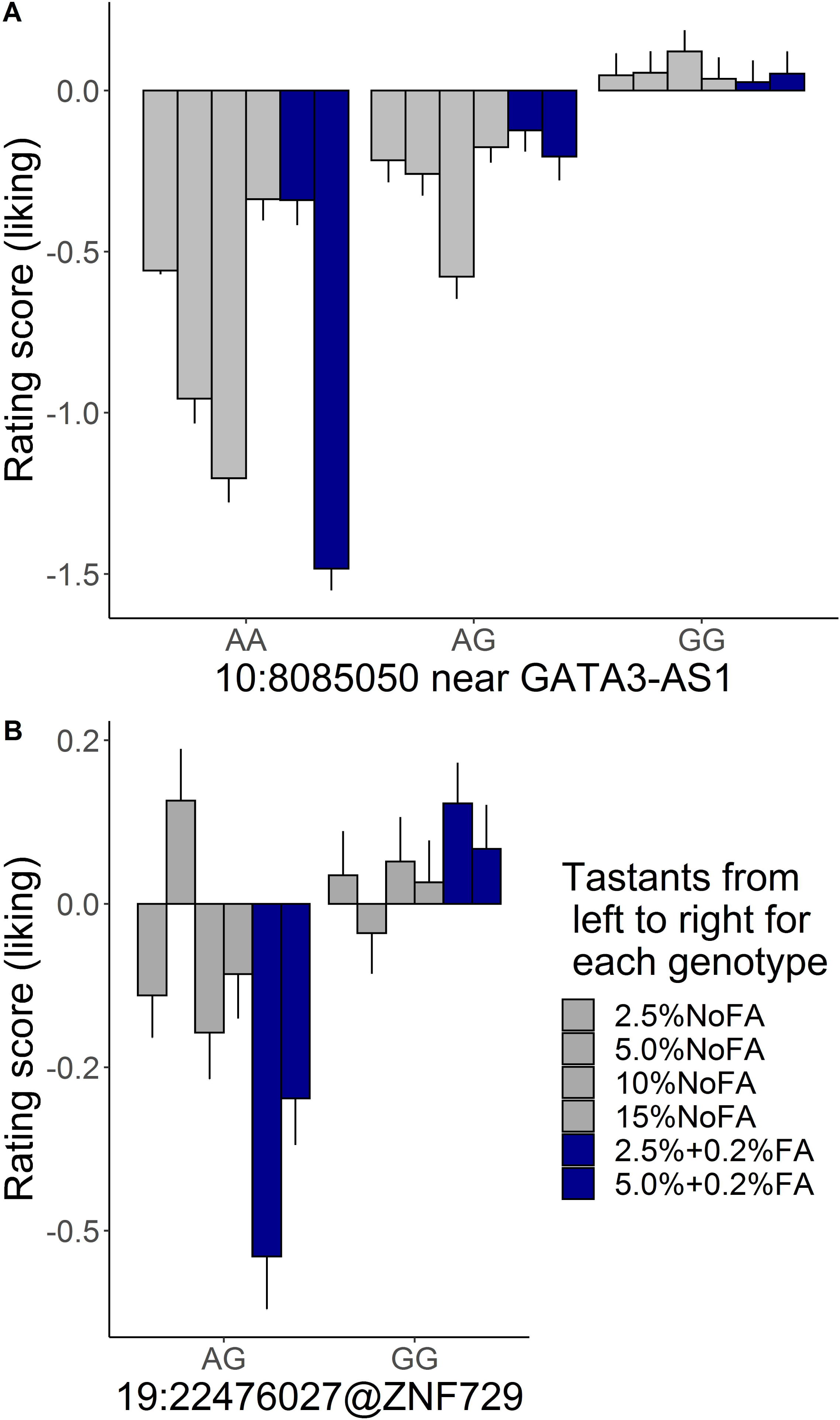
Allele effect of variants *10:8085050* near gene *GATA3-AS1* (**A**) and *19:22476027* within the gene *ZNF729* (**B**) on ratings of liking across types of potato chips. For both variants, participants with G allele rated higher liking for all potato chip types than did those with other allele. The standard residual scores for liking were calculated in the general linear model with covariates of sex, age, and 20 eigenvalues.

**Table 3.** Suggestive associations for ratings of potato chip fattiness and liking identified by uvGWAS.

| Stimuli | CHR | SNP (CHR:BP) | ALT | REF | MAF | Beta | SE | P | Gene | HIT_TYPE | SNP_TYPE |
| --- | --- | --- | --- | --- | --- | --- | --- | --- | --- | --- | --- |
| **Fattiness** |  |  |  |  |  |  |  |  |  |  |  |
| 5.0% NoFA | 4 | 4:45361270 | G | C | 0.12 | 0.92 | 0.20 | 6.14E-06 | *PRKRIRP9* | nearest | Imputed |
| 5.0% NoFA | 7 | 7:42032565 | T | C | 0.26 | 0.62 | 0.14 | 8.25E-06 | *GLI3* | within | Genotyped |
| 5.0% NoFA | 8 | 8:105135473 | A | T | 0.15 | 0.86 | 0.18 | 1.92E-06 | *RIMS2* | within | Imputed |
| 5.0% NoFA | 11 | 11:7476601 | A | G | 0.21 | 0.74 | 0.15 | 9.51E-07 | *SYT9* | within | Imputed |
| 10% NoFA | 7 | 7:25514020 | C | G | 0.18 | 0.74 | 0.16 | 4.91E-06 | *AC091705.1* | nearest | Imputed |
| 15% NoFA | 5 | 5:94138997 | C | T | 0.17 | 0.76 | 0.17 | 4.37E-06 | *MCTP1* | within | Genotyped |
| 15% NoFA | 6 | 6:77345798 | C | T | 0.10 | 0.87 | 0.20 | 9.64E-06 | *RP11-354K4.1* | nearest | Imputed |
| 15% NoFA | 11 | 11:86707909 | G | C | 0.09 | 0.88 | 0.20 | 9.45E-06 | *RP11-736K20.6* | within | Imputed |
| 15% NoFA | 19 | 19:51226244 | T | C | 0.38 | 0.59 | 0.13 | 9.19E-06 | *CLEC11A* | nearest | Imputed |
| **Liking** |  |  |  |  |  |  |  |  |  |  |  |
| 5.0% NoFA | 2 | 2:215358759 | C | T | 0.06 | -1.12 | 0.25 | 8.47E-06 | *VWC2L* | within | Imputed |
| 5.0% NoFA | 20 | 20:50443885 | T | C | 0.25 | 0.63 | 0.13 | 1.84E-06 | *RP5-1112F19.2* | nearest | Imputed |
| **10% NoFA** | **10** | **10:8085050** | **A** | **G** | **0.08** | **-0.92** | **0.20** | **5.27E-06** | **GATA3-AS1** | **nearest** | **Imputed** |
| 10% NoFA | 12 | 12:127482374 | C | G | 0.07 | -0.95 | 0.21 | 9.08E-06 | *RP11-575F12.1* | within | Imputed |
| 2.5% with 0.2% FA | 4 | 4:160267304 | A | G | 0.06 | -1.32 | 0.30 | 8.02E-06 | *RAPGEF2* | within | Imputed |
| 2.5% with 0.2% FA | 12 | 12:13948270 | A | G | 0.10 | -1.13 | 0.24 | 3.58E-06 | *GRIN2B* | within | Imputed |
| **2.5% with 0.2% FA** | **19** | **19:22476027** | **A** | **G** | **0.11** | **-1.07** | **0.23** | **4.92E-06** | **ZNF729** | **within** | **Imputed** |
| 5.0% with 0.2% FA | 13 | 13:95498608 | T | C | 0.09 | -0.98 | 0.22 | 5.94E-06 | *RPL21P112* | nearest | Imputed |

**Table 4.** Suggestive genes for ratings of potato chip fattiness and liking identified by mvGWAS

| Trait | CHR | SNP<br>(CHR:BP) | ALT | REF | MAF | P | Gene | HIT_TYPE | SNP_TYPE |
| --- | --- | --- | --- | --- | --- | --- | --- | --- | --- |
| Fattiness | 2 | 2:141755773 | A | T | 0.32 | 2.96E-06 | *LRP1B* | within | Imputed |
|  | 10 | 10:116662107 | C | T | 0.05 | 8.97E-06 | *RP11-106M7.4* | nearest | Imputed |
| Liking | 7 | 7:34676953 | T | A | 0.06 | 3.05E-07 | *NPSR1-AS1* | within | Imputed |
|  | 8 | 8:40998854 | A | T | 0.04 | 5.14E-07 | *RNU6-356P* | nearest | Imputed |
|  | 9 | 9:20636973 | T | C | 0.48 | 1.15E-06 | *MLLT3* | nearest | Genotyped |
|  | **10** | **10:8085050** | **G** | **A** | **0.08** | **8.19E-07** | **GATA3-AS1** | **nearest** | **Imputed** |
|  | **19** | **19:22476027** | **A** | **G** | **0.11** | **4.71E-06** | **ZNF729** | **within** | **Imputed** |

### Candidate genes

None of the candidate genes consistently met a genome-wide statistical threshold, but some candidate genes were more often associated with potato chip fattiness or liking than others at a nominal significance threshold (*p* < 0.05; **Figure 6A**). The most notable results were significant variants within *CD36* and *TRPM5* associated with potato chop liking and fattiness (**Figure 6B, C**; **Supplemental Figure 8**, **Supplemental Tables 3 and 4**). For *CD36*, the variant *rs1761667* (which was associated with fat perception in previous studies) did not pass the quality-control filters, but we examined a nearby variant, *rs1722501,* that was in nearly perfect linkage disequilibrium (*R*^2^ > 0.99) with *rs1761667*. However, participants did not differ in ratings of potato chip fattiness or liking for this proxy marker (**Supplemental Table 5**), although there were many associations for other variants within *CD36*, as noted above (see **Figure 6**).

**Figure 6.**
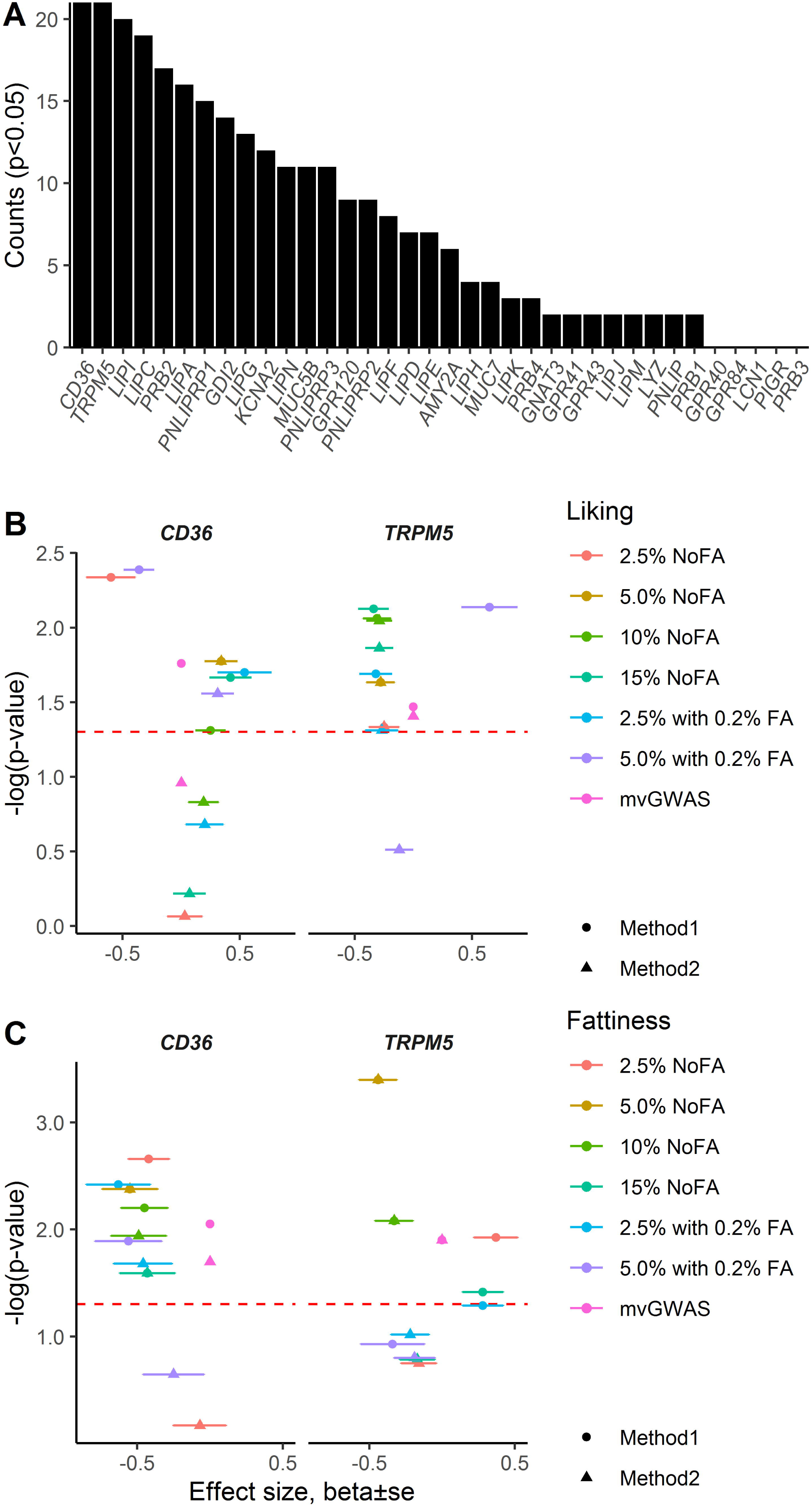
Candidate gene effect on fat perception for potato chips. (**A**) Total counts of nominal *p* < 0.05 out of 28 tests for each candidate gene for the two methods of candidate gene analysis (method 1 and method 2; see Materials and Methods) in the outputs from uvGWAS and mvGWAS. (**B, C**) Associations of top variants within candidate genes *CD36* and *TRPM5* with ratings of liking (**B**) and fattiness (**C**) for each type of potato chip. x-Axes show effect size (β±SE), obtained from uvGWAS, and y-axes show –log(*p*-value), obtained from uvGWAS and mvGWAS, for the top variants within ±SE data were available from mvGWAS; i.e., β±SE=0 is not true). Red dashed lines indicate *p* = 0.05; the points above this line indicate a nominal significant effect on the trait. FA=fatty acid (hexadecanoic acid). For other details of the data, see **Supplemental Tables 3 and 4**.

### Gene expression, pathway, and gene enrichment analysis

We reasoned that expression of fat candidate genes (those that have a proposed role in peripheral fat or fatty acid signaling) would be a benchmark to compare the taste-tissue expression of the novel genes identified from the GWAS results. Compared with receptor and other signaling candidate genes (*GPR40*, *GPR41*, *GRP43*, *GPR84*, *GPR120*, *TRPM5*, *CD36*, *KCNA2*, and *GNAT3*), the novel genes have relatively higher expression levels in fungiform papillae, especially for *RAPGEF2, GLI3, MCTP1*, and *MLLT3* (**Figure 7**). *ZNF729* and *GATA3-AS1* had a similar expression abundance as the candidate genes *GPR40*, *GPR41*, *GPR84*, *GPR120*, *KCNA2*, and *TRPM5* but much lower than the candidate genes *GRP43*, *CD36*, and *GNAT3*. The presence of many of the novel genes in taste tissue is consistent with a role in peripheral perception, but some candidate genes had a very low abundance. This subset of low-abundance novel genes may be nearly undetectable in the taste tissue sampled because only a few of the relevant cells may have been present in the tissue sample or because the genes may act at different times (e.g., early development) or in different tissues (e.g., the filiform papillae or the brain).

**Figure 7.**
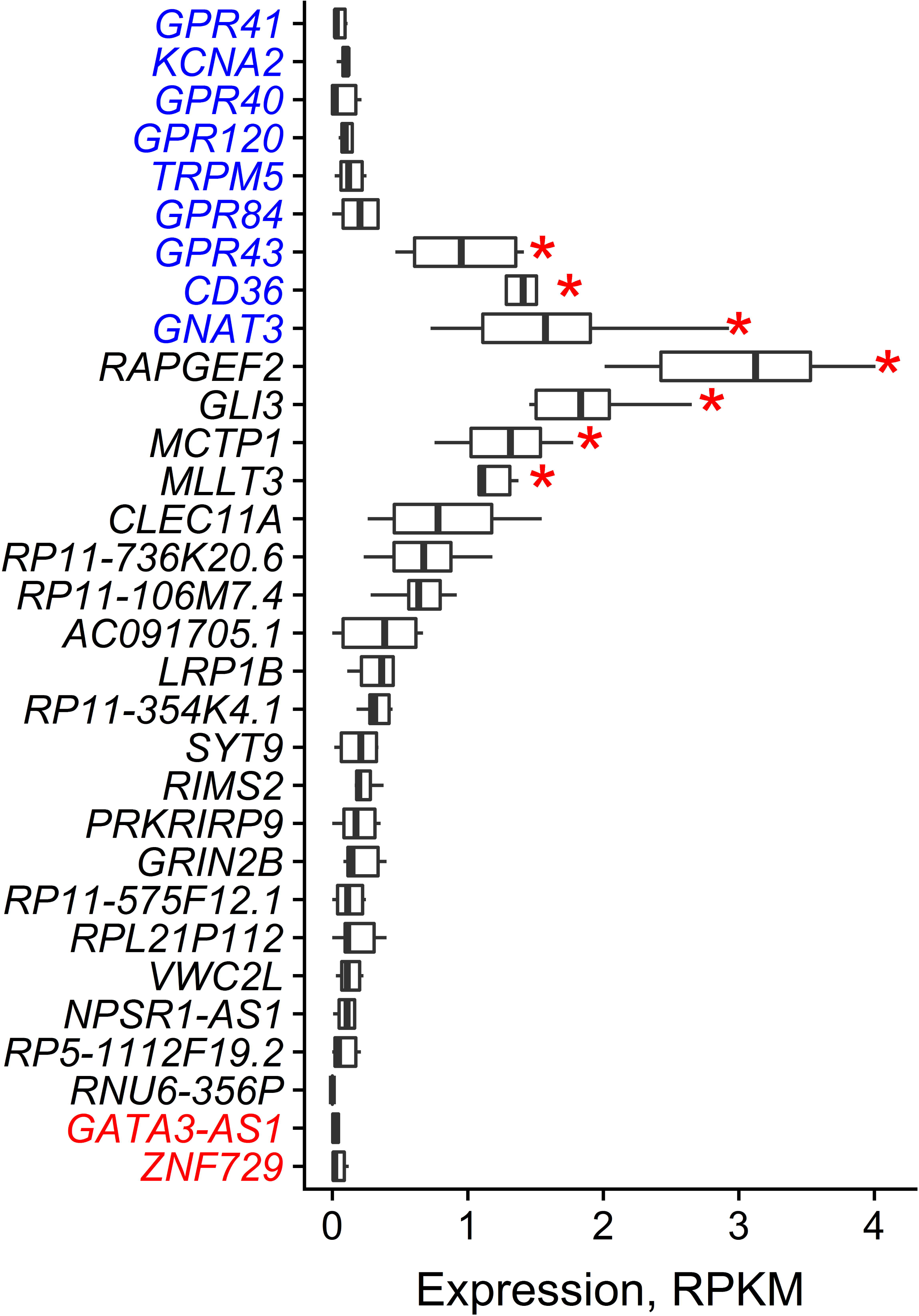
Box plots of taste tissue expression abundance of genes near the peak statistical associations from the GWAS (novel hits) and for candidate genes (shown in blue) known from prior studies to contribute to fat perception. Two genes, *ZNF729* and *GATA3-AS1* (shown in red), were commonly detected by both uvGWAS and mvGWAS in the present study. RPKM=reads per kilo base per million mapped reads. *RNU6-356P* had no expression in any sample. Outliers are not shown. Red asterisks indicate genes with statistically higher expression level compared with other genes in taste tissue (*p* < 0.05/351 = 0.000142, Bonferroni corrections for multiple tests).

We conducted pathway analysis to understand the function of as many of the novel genes identified as possible. In the GENEVESTIGATOR analysis, 21 of the 22 associated genes identified by GWAS (*RP11-575F12.1* is not found in the database) were tested against the 74,727 background genes. Three gene sets were enriched using the associated genes as bait (*p* < 0.001, Fisher’s exact test; **Supplemental Figure 9**, **Supplemental Table 6**), from the Gene Ontology categories synapse GO:0045202, cell-cell signaling GO:0007267, and positive regulation of neurogenesis GO:0050769. Overall, these results point to a role of these genes and their protein products in sensory signaling and perhaps regulation of sensory cell types.

## Discussion

Dietary fat is added to food to increase its flavor and palatability, but whether fat is sensed by chemical cues (e.g., from fatty acids), textural cues, or both is contentious. The data from this study support previous observations that fatty acids provide a chemical cue for fattiness but that this component of fattiness is not desirable (Running *et al.* 2017). When hexadecanoic acid (a saturated 16-carbon fatty acid) was added to the potato chip lowest in fat, it was rated as fattier but was less liked than a potato chip with a comparable amount of fat but without the added fatty acid. Thus, presumably, taking a broader view and generalizing, this result suggests that increasing “fattiness” by adding fatty acids to foods would not make them better liked, and raises the possibility that recently discovered antagonists to the fatty acid receptors (Milligan *et al.* 2017) might improve fat flavor. These data support the hypothesis that there are at least two sensory inputs for fat perception, a chemical cue and presumably a textural cue, with the texture conveying perhaps the pleasant aspects of fattiness.

In addition to studying the relationship between fattiness and liking, we also attempted to study fat discrimination, asking participants to choose the fattier milk solution from a pair of high- and low-fat samples. This task was difficult for the participants, and almost no one correctly identified the high-fat sample 10 times out of the 10 trials. This result came as a surprise because our preliminary testing suggested this task was easy; however, most preliminary testing was conducted with commercially available low- and high-fat milk samples and in a quiety sensory laboratory, making discrimination easier. The prepared milk samples used for testing here were the same in all aspects except the amount dietary fat added, and for many people the oral cues alone (as opposed to visual or olfactory cues) are insufficient to discriminate low-fat from high-fat samples. One additional concern about data was the effect of transportation on the stimuli: the milk was prepared and then driven by truck several hundred miles to the test location – conceivably, vibration may have caused coalescence of the fat globules that altered the ability to discriminate between samples.

The main focus of this study was to examine whether person-to-person differences in the liking or perception of fattiness are due in part to individual genetic variation. To establish the heritability of a trait, it is essential to have a reliable measurement, that is, a trait that can be measured reproducibly; accordingly, demonstrating that the measures used were reliable was an essential precondition for the heritability calculations. We learned from the reliability and heritability analyses that liking for this solid food matrix, potato chips, with differing fat concentrations was more similar among genetically identical (MZ) twins than among nonidentical (DZ) twins. Ratings of fattiness were also heritable, but less so, aligning with results from our studies of other taste modalities, which, for example, demonstrated that liking for a concentrated salt solution is more heritable than are salty intensity ratings (Knaapila *et al.* 2012). Our results are in contrast to a recent study of the effect of diet on fatty acid perception in twins, which reported few or no genetic effects (Costanzo *et al.* 2018); however, these two studies differed in methods, as did the number of twins investigated, 88 in (Costanzo *et al.* 2018) vs. 398 here.

Thus, despite the logistical challenges posed by measuring percepts from dietary fat, there is evidence for a genetic determinant on par with other traits that have been studied using GWAS methods (Clarke *et al.* 2017). Building on the heritability analysis, we also performed two types of GWAS, which are agnostic to prior information about which genes and variants might be previously known or suspected to contribute to the perception of dietary fat. This part of the study was underpowered and returned no results that met the classic statistical threshold for GWAS results but did provide, in tandem with the bioinformatic analysis, clues about which genes and pathways might be worth pursuing in future work, specifically in the realm of cell-to-cell communication and perhaps cell type.

Of particular interest is the association between fat liking and variants in the transcription factors that contribute to the development of taste cells (Ermilov *et al.* 2016; Qin *et al.* 2018). The transcriptome was not helpful in interpreting the novel genes in part because taste tissue from fungiform papillae is unlikely to be involved in the textural aspects of fat perception, and in part because the abundance of even the known genes is very low to undetectable in fungiform taste tissue. Single-cell studies from all regions of the oral cavity would be a step forward, which is increasingly more feasible as methods improve, although the most complete experimental paradigm would also include the sensory pathways, including brain regions that process the sensory properties of dietary fat information (Grabenhorst and Rolls 2014).

The results of the candidate gene analyses were more compelling in the sense that, although none of the results were individually very striking, multiple methods of analysis have repeatedly indicated a role for *CD36* and *TRPM5* in the perception of dietary fat, in both human and animal studies (Chamoun *et al.* 2018), especially gene knockout studies. Parenthetically, we did not see associations with the proxy marker we used to try to replicate the previous studies exactly (Keller *et al.* 2012; Mrizak *et al.* 2015; Pepino *et al.* 2012; Sayed *et al.* 2015), but *CD36* is a large gene with many potentially functional variants, and therefore a fine-mapping study in multiple populations is warranted. There may be multiple variants that cause a spectrum of effects that differ by ancestral population, e.g., (Gurdasani *et al.* 2019).

We speculate that sensory nutrition and taste perception offer a way to reduce nutrition-related human diseases, by studying the nuanced and often misunderstood relationship between liking and intake (Hayes in press). GWAS allows us to screen and identify common genetic variants associated with fat consumption (Tanaka *et al.* 2013), and our findings, combined with future functional genomic analyses, especial single-cell profiling, will delineate the causal genetic variants and biological mechanisms underlying the observed statistical associations with fat perception (Gallagher and Chen-Plotkin 2018).

## Acknowledgments

We thank the following people for assistance with data collection, listed in alphabetical order: Charles J. Arayata, Nuala Bobowski, Fujiko Duke, Hillary Ellis, Brad Fesi, Nicole Greenbaum, Aurora Hannikainen, Desmond Johnson, Katherine Leung, Durpri Lin, Alex Mangroo, Corrine Mansfield, Michael Marquis, Elliott McDowell, Tiffany Murray, Lauren Shaw, Lindsey Snyder, Molly Spencer, Amber Suk, Alyssa Treff, and Casey Trimmer. We thank the twins for their participation and the administration of TwinsDays including Sandy Miller and Janine Bregitzer for their assistance during data collection.

---

**Supplemental Figure 1.** Changes in ratings of fattiness and liking by corn oil concentration across the six types of potato chips. FA=fatty acid (hexadecanoic acid). For other details, see **Figure 1**.

**Supplemental Figure 2.** Violin plots for ratings of the sensory traits. The violin area shows the estimated density of each rating score point. The dots and bars show means and SDs.

**Supplemental Figure 3.** Pearson correlations between test and retest of each rating (*n* = 50).

**Supplemental Figure 4.** Pearson correlations between age and sensory measures for potato chips and other taste stimuli. Young participants were more sensitive to taste stimuli than were older participants.

**Supplemental Figure 5.** Least square mean (LSM) and standard error of sensory measures by race. EA=European Americans, AA=African Americans, Oth=others (Asian, Hispanic, Native American, mixed). Different letters (a, b) show a significant LSM difference.

**Supplemental Figure 6.** Most participants had difficultly discriminating milk fat content, with near chance levels overall. (**A**) Histogram of milk fat discrimination scores. The dashed white line shows probability of success, which is near the chance level of 5, but it is significantly different from the chance level, *p* < 0.001. (**B**) No significant correlations were observed between twin 1 and twin 2 for milk discrimination for either DZ or MZ twins; thus, no heritability for milk fat discrimination scores was calculated.

**Supplemental Figure 7.** Regional associational plots, based on mvGWAS results, for single-nucleotide polymorphisms in linkage disequilibrium (*r*^2^) with the peak variants *10:8085050* near the gene *GATA3-AS1* (**A**) and *19:22476027* within the gene *ZNF729* (**B**) for ratings of liking, and for the fat perception candidate gene *CD36* for ratings of liking (**C**) and fattiness (**D**) for potato chips. The highlighted chromosome regions show the target genes.

**Supplemental Figure 8.** Associations of top variants within each candidate gene with ratings of liking (A) and fattiness (B) for each type of potato chip. For details see **Figure 6**.

**Supplemental Figure 9.** Gene set enrichment analyses. Venn diagram visualizes overlapping genes among the top three gene sets and the target genes (21 out of 22 GWAS hits; *RP11-575F12.1* is not found the database). All genes (*n* = 74,727 total genes) were selected from Reactome annotations and Gene Ontology annotations as background collection. The top three gene sets identified are synapse GO:0045202, cell-cell signaling GO:0007267, and positive regulation of neurogenesis GO:0050769 (see **Supplemental Table 6**).

**Supplemental Table 1. Summary statistics for linear mixed model analyses**

FA, fatty acid; ICC, intraclass correlation. Boldface indicates the test statistic meets a significance threshold of *p* < 0.01.

**Supplemental Table 2.** The effect of sex, race, and age on sensory measures for potato chips, taste stimuli, and milk discrimination

PTC, phenylthiocarbamide. Highlighting indicates suggestive effects with a *p-value* < 0.05.

**Supplemental Table 3**. The effect of the top variant within each candidate gene on ratings of potato chip fattiness and liking.

FA, fatty acid; mvGWAS, multivariate genome-wide association study. Highlighting indicates suggestive effects with a *p-value* < 0.05.

*For *GPR41* and *GPR84*, no variant within the genes was available from the association data, so we expanded the region to 500 bp up- and downstream for each site when extracting the variant to examine for association. For other details see **Supplementary Tables 5 and 6**.

**Supplemental Table 4.** The top variant within each candidate gene with effects on ratings of potato chips with 5% corn oil (without added fatty acid) had effects on fattiness and liking for other types of potato chips. For details, see **Supplemental Table 3**.

**Supplemental Table 5.** The variant *rs1722501* (chr7:80244694) as proxy for *rs1761667* within *CD36* has no significant effect on ratings of potato chip fattiness and liking.

FA, fatty acid; mvGWAS, multivariate genome-wide association study. *rs1761667* (7:80244939) was not in Hardy-Weinberg disequilibrium (*p*=8.5e-15) and thus did not pass the filter test statistics (*p*>1e-6); therefore, we extracted the variant *rs1722501*, which had an *R*^2^>0.99 and linkage disequilibrium>0.99 with *rs1761667*. For this marker, there was no significant effect on fatty and liking for any of the six potato chip types tested. For abbreviations, e.g., MAF, see **Tables 3**.

**Supplemental Table 6.** Gene set enrichment analysis

